# TRF2 rescues telomere attrition and prolongs cell survival in Duchenne muscular dystrophy cardiomyocytes derived from human iPSCs

**DOI:** 10.1101/2022.01.10.475653

**Authors:** Asuka Eguchi, Sofía I. Torres-Bigio, Kassie Koleckar, Adriana Fernanda G. S. Gonzalez, Foster Birnbaum, Helen M. Blau

## Abstract

Duchenne muscular dystrophy (DMD) is a severe muscle wasting disease caused by the lack of dystrophin. Heart failure, driven by cardiomyocyte death, fibrosis, and the development of dilated cardiomyopathy, is the leading cause of death in DMD patients. Current treatments decrease the mechanical load on the heart, but do not address the root cause of dilated cardiomyopathy: cardiomyocyte death. Previously, we showed that telomere shortening is a hallmark of cardiomyocytes in DMD patient cardiac tissues and in a mouse model of DMD we generated that manifests dilated cardiomyopathy. We also found that this telomere shortening is recapitulated in DMD cardiomyocytes differentiated from patient-derived induced pluripotent stem cells (iPSCs). Here we further characterize that DMD cardiomyocytes exhibit reduced cell size, nuclear size, and sarcomere density compared to healthy isogenic controls. The telomere-binding protein, TRF2, is a core component of the shelterin complex, which protects chromosome ends and is expressed at lower levels in DMD cardiomyocytes compared to controls. We investigated whether preservation of telomere length could be achieved with upregulation of TRF2. We found that TRF2 upregulation in DMD cardiomyocytes increased telomere lengths, cell size, nuclear size, sarcomere density, and cell survival. These data suggest TRF2 gene therapy has the potential to delay the onset of dilated cardiomyopathy.

## Introduction

Duchenne muscular dystrophy (DMD) affects 1 out of 5,000 males and manifests as progressive muscle degeneration (1). DMD is caused mutations in dystrophin, which has an essential role in connecting the contractile apparatus to the extracellular matrix and is encoded by the largest gene in the genome (2, 3). Dilated cardiomyopathy is the leading cause of death in DMD patients (4). Currently, patients are prescribed angiotensin-converting enzyme inhibitors, angiotensin II receptor blockers, and mineralocorticoid receptor antagonists that lower blood pressure or beta blockers that slow the heart rate (5). These strategies reduce the mechanical load on the heart, but do not address the cardiomyocyte death and fibrosis that drive the progression of dilated cardiomyopathy.

We previously reported an intriguing link between genetic cardiomyopathies and telomere shortening. The dystrophin-deficient *mdx* mouse does not manifest the characteristic cardiac symptoms of DMD. However, when crossed with an *mTR* knock-out mouse that is missing the RNA component of telomerase (6, 7), *mdx*^4cv^/*mTR*^*G*2^ mice exhibit hallmark DMD phenotypes, including severe muscle degeneration, kyphosis, dilated cardiomyopathy, and shortened lifespan (8–10). Notably, in the hearts of these mice, the telomeres are shortened specifically in the cardiomyocytes but not in the smooth muscle cells or spermatocytes indicating that the effect is not systemic (9).

For unknown reasons, mice have significantly longer telomeres than humans (11). Our discovery that *mdx*^4*cv*^/*mTR*^*G*2^ but not *mdx* mice develop dilated cardiomyopathy, suggested that long telomeres protect them from the cardiac failure characteristic of human DMD (9). In corroboration, we found that telomere shortening is a common feature of cardiomyocytes from tissues of patients with cardiomyopathies caused by mutations in genes that encode sarcomeric or cytoskeletal proteins, including dystrophin (12). Telomere lengths in DMD cardiomyocytes were shorter than those of age-matched healthy controls (12).

Furthermore, mutations in *Notch1* lead to premature calcification of the aortic valve (13), yet mice with the same heterozygous for *Notch1* mutation do not exhibit this pathogenic phenotype until they are bred with mTR^G2^ knock-out mice and their telomeres are shortened or “humanized.” These findings underscore the role of long telomeres as being cardioprotective (14).

We and others have capitalized on human induced pluripotent stem cells (iPSCs) to model cardiac disease phenotypes in a dish in order to surmount differences between human and mouse cells (12, 15–17). We have shown that in cardiomyocytes differentiated from iPSCs (iPSC-CMs) derived from patients harboring mutations in contractile proteins including dystrophin, titin, and troponin T (12), telomeres shorten in conjunction with the development of cardiomyocyte deficits (15). Such shortening does not occur in isogenic control iPSC-CMs in which the mutations are CRISPR-corrected (12, 15). Here, we investigate whether telomere shortening can be abrogated and whether telomere protection ameliorates the pathogenic characteristics of DMD cardiomyocytes. We use three DMD iPSC lines with matching healthy control lines and at least three differentiation experiments per line, a robust dataset. By using isogenic controls, we ensure we control for genetic background and that the differences we observe between healthy and diseased states can be attributed to the dystrophin deficiency. We recently reported that DMD iPSC-CMs exhibit aberrant calcium handling, impaired contractile function, and shortened telomeres, leading to a DNA damage response (15). Here we extend these studies on DMD iPSC-CMs as a disease model by showing that the telomere shortening, previously demonstrated by quantitative fluorescent in situ hybridization (Q-FISH), is also observed by Southern blots of telomere restriction fragments. We capitalize on a lentiviral troponin T reporter system with a zeocin resistance gene to enrich for cardiomyocytes. We also observe morphological deficits in DMD iPSC-CMs that inform the contractile deficits we have previously reported. We test whether the homodimeric shelterin protein, TRF2, can protect the telomeres from shortening. We find that TRF2 prevents telomere shortening and rescues pathogenic characteristics of smaller cell size, reduced sarcomere density, and premature cell death in DMD iPSC-CMs. We also capitalize on a bioengineered platform to micropattern iPSC-CMs to a physiological length:width aspect ratio for calcium imaging. Our data highlight a therapeutic strategy, increased expression of TRF2, to ameliorate pathogenic characteristics of DMD cardiomyocytes. Although not a cure, in contrast to gene correction or exon skipping strategies, our therapeutic strategy has the potential to universally address all DMD mutations.

## Results

### DMD iPSC as a Disease Model

We differentiated human induced pluripotent stem cells (iPSCs) to cardiomyocytes to model Duchenne muscular dystrophy in culture (Table 1 and Fig. 1*A*). Dystrophin, the protein that is missing in DMD, is encoded by the largest gene in the human genome and encompasses 2.4 megabases (18). As a result, DMD can be caused by a wide variety of mutations that impact the function of the dystrophin protein (19). Two lines, DMD19 and DMD16, were derived from patients with nonsense mutations in the dystrophin gene that result in degradation of the transcript by nonsense-mediated mRNA decay (20, 21). Both lines have been CRISPR-corrected to generate corresponding isogenic controls, DMD19 iso and DMD16 iso, in which the dystrophin mutations are corrected (20, 21). A third line, UC3.4, was derived from a healthy patient, and the isogenic DMD line, UC1015.6, was CRISPR-induced to yield a DMD mutation, c.263delG, which results in deletion of the N-terminus of dystrophin that renders the truncated protein dysfunctional (22, 23). Importantly, the lack of the N-terminal actin-binding domain of dystrophin is associated with early onset dilated cardiomyopathy (24). The use of three DMD iPSC lines with matching isogenic healthy controls ensures that results are controlled for genetic background, and consistent differences observed between healthy and diseased states can be attributed to the dystrophin deficiency.

**Table 1.** iPSC lines and mutational status.

| Category | Name | Mutation Type | Description | Reference |
| --- | --- | --- | --- | --- |
| Control | DMD19 iso | None | Isogenic control of DMD19 | (20, 21) |
| DMD | DMD19 | Patient-derived | DMD (c.4918_4919delACinsTG) | (20, 21) |
| Control | DMD16 iso | None | Isogenic control of DMD16 | (20, 21) |
| DMD | DMD16 | Patient-derived | DMD (c.10171C>T) | (20, 21) |
| Control | UC3.4 | None | Isogenic control of UC1015.6 | (22, 23) |
| DMD | UC1015.6 | CRISPR-induced | DMD (c.263delG) | (22, 23) |

**Figure 1.**
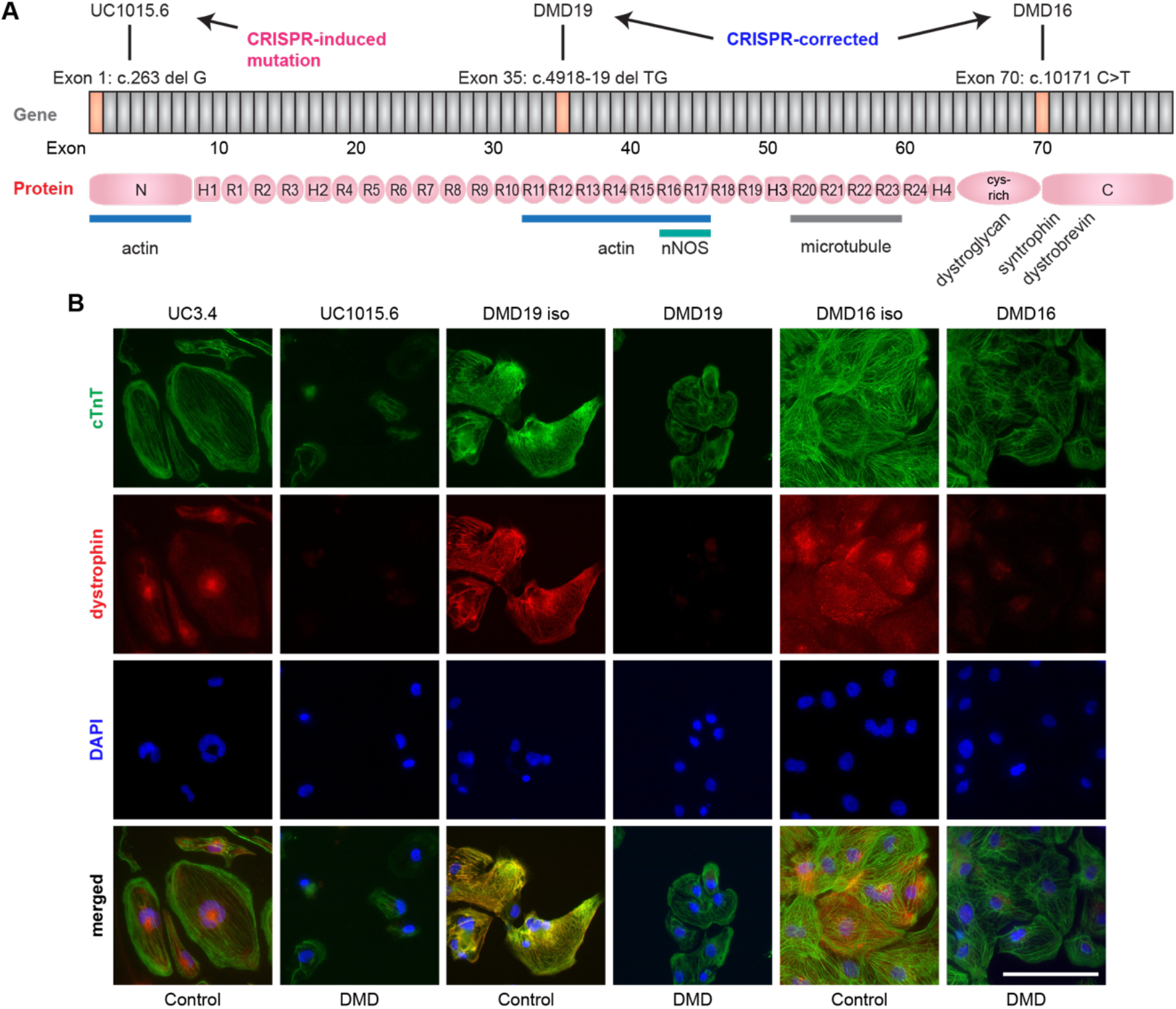
Cardiomyocytes differentiated from dystrophin-deficient induced pluripotent stem cells. (*A*) Schematic of dystrophin gene and the mutations corresponding to the cell lines. The UC1015.6 line harbors a CRISPR-induced mutation that results in expression of a truncated dystrophin missing the N-terminus. DMD19 and DMD16 are patient-derived iPSCs that have nonsense mutations. (*B*) Cardiac troponin T (cTnT) and dystrophin immunostaining in Day 30 iPSC-CMs. The UC lines were stained with the MANEX1A antibody that recognizes the N-terminus of dystrophin. The DMD19 and DMD16 lines were stained with the ab15277 antibody that recognizes the C-terminus. DAPI in blue marks nuclei. Scale bar = 100 μm.

All six iPSC lines expressed pluripotency markers, OCT4, SOX2, and TRA-1-60, before differentiation (*SI Appendix*, Fig. S1*A-C*). We differentiated the iPSCs to cardiomyocytes using established protocols (25), then purified the iPSC-CMs by culturing them in media lacking glucose and supplemented with lactate, a method that starves contaminating fibroblasts (26). On Day 30, we observed cardiac troponin T (cTnT) immunostaining in the iPSC-CMs (Fig. 1*B*). We also immunostained iPSC-CMs for dystrophin and observed expression along the sarcomeres in the healthy control lines, UC3.4, DMD19 iso, and DMD16 iso (Fig. 1*B*). As expected, the DMD lines, DMD19 and DMD16, did not show dystrophin expression along the sarcomeres as they harbor nonsense mutations (Fig. 1*B*) (21). In contrast, the UC1015.6 line has a mutation that induces the expression of a truncated dystrophin lacking a large portion of the N-terminal domain that binds actin filaments (23). Immunostaining with the MANEX1A antibody that recognizes the N-terminus of dystrophin encoded by exon 1 showed a lack of expression in the DMD line, UC1015.6 (Fig. 1*B*) (27), and use of the ab15277 antibody that recognizes the C-terminus detected the truncated dystrophin (*SI Appendix*, Fig. S2*A*), consistent with the report that this line expresses a dystrophin variant missing the N-terminal actin-binding domain (23). All lines, healthy and DMD alike, showed nuclear staining with the C-terminal antibody. The ubiquitously expressed Dp71 isoform, expressed in human heart tissue and in iPSC-CMs (21,28, 29), is the protein form likely recognized as none of the mutations we use here preclude expression of this isoform. Taken together, our characterization of the iPSC-CMs demonstrated expression of cardiac-specific markers and muscle-specific dystrophin deficiency in the diseased cells and validated use of these lines as robust disease models to study the role of telomeres in the etiology of DMD.

### DMD iPSC-CMs Exhibit Morphological Deficits

We investigated the morphological deficits that could potentially account for functional shortcomings. Previously, we reported that DMD iPSC-CMs exhibit impaired contractile function on stiff substrates mimicking a fibrotic myocardium as well as poor calcium handling (15). On Day 30 of differentiation, we observed that DMD iPSC-CMs were smaller in cell size and had smaller nuclei (Fig. *2A-I*). The reduced cell size is consistent with our previous report where we seeded iPSC-CMs on 2,000 μm^2^ micropatterns of extracellular matrix with a length:width aspect ratio of 7:1 (15), a regimen that has been shown to promote iPSC-CM maturation (30–32). As shown here, when cultured in monolayers without any physical constraints, the difference in cell sizes were more pronounced. DMD iPSC-CMs were on average 58% the size of the healthy iPSC-CMs (Fig. 2*D-F*). Strikingly, in *mdx*^4*cv*^/*mTR*^*G*2^ mice, we also observed reduced cardiomyocyte diameter and smaller nuclei in the hearts of the mouse model that phenocopies the severe symptoms observed in Duchenne patients (9). Morphological changes to the nuclei have also been reported in histological characterization of cardiac tissue from DMD patients with signs of nuclear pyknosis, where the chromatin condenses during cell death (33, 34).

**Figure 2.**
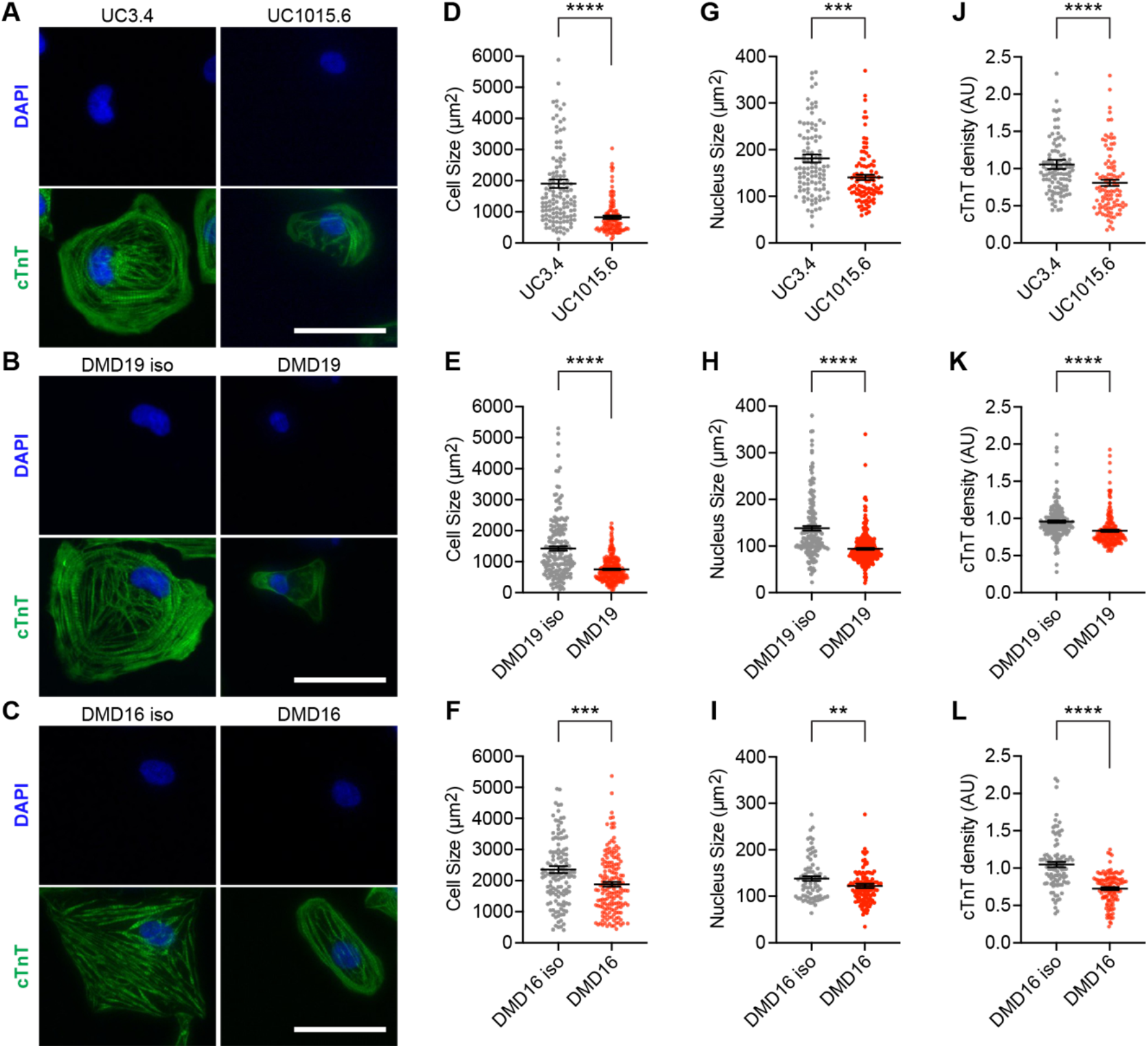
DMD iPSC-CMs exhibit deficits in cell size, nuclear size, and sarcomere density on Day 30 of differentiation. Cardiac troponin T (cTnT) immunostaining and DAPI staining for nuclei in (*A*) UC3.4 and UC1015.6 iPSC-CMs, (*B*) DMD19 iso and DMD19 iPSC-CMs, and (*C*) DMD16 iso and DMD16 iPSC-CMs. Scale bar = 50 μm. Area of cells for (*D*) UC3.4 and UC1015.6 iPSC-CMs, (*E*) DMD19 iso and DMD19 iPSC-CMs, and (*F*) DMD16 iso and DMD16 iPSC-CMs. Nuclear size for (*G*) UC3.4 and UC1015.6 iPSC-CMs, (*H*) DMD19 iso and DMD19 iPSC-CMs, and (*I*) DMD16 iso and DMD16 iPSC-CMs. Sarcomere density as measured by cTnT signal over cell area for (*J*) UC3.4 and UC1015.6 iPSC-CMs, (*K*) DMD19 iso and DMD19 iPSC-CMs, and (*L*) DMD16 iso and DMD16 iPSC-CMs. Cells were scored from 3 differentiation experiments. N = 97-205 cells. Data shown are mean ± SEM. One-way ANOVA and Mann-Whitney test for post-hoc comparison was used to determine significance. * p < 0.05, ** p < 0.01, *** p < 0.001, and **** p < 0.0001.

The sarcomere density as measured by cTnT signal over cell area was also reduced in DMD iPSC-CMs compared to isogenic controls (Fig. *2J-L*). This decrease in cTnT levels was independent of cell size, indicating that DMD iPSC-CMs are both smaller and have less sarcomeres. The reduced sarcomere content likely accounts for the reduced force of contraction, contraction velocity, and relaxation velocity, which we previously measured in these DMD iPSC-CMs (15). Importantly, our findings are consistent with electron micrographs that showed sparse myofibril content in the hearts of DMD patients (33, 34). The reduced cell size, nuclear size, and sarcomere density were statistically significant in all three lines of DMD iPSC-CMs when compared with the healthy isogenic controls, underscoring the significance of these results. Collectively, the morphological deficits of DMD iPSC-CMs suggest that their smaller cell size as well as reduced sarcomere content impairs their ability to contract like healthy cells. TRF2 Rescues Telomere Attrition in DMD iPSC-CMs. Since we previously determined that telomeres are shortened in DMD iPSC-CMs (12, 15), we hypothesized that a key shelterin protein, telomeric repeat-binding factor 2 (TRF2), might be expressed at lower levels. We measured TRF2 levels by western blot and found that DMD iPSC-CMs express lower levels of TRF2 compared to isogenic controls (*SI Appendix*, Fig. S2*E*). We sought to test whether overexpression of TRF2 could rescue telomere attrition in DMD iPSC-CMs. In addition to enrichment of the cardiomyocyte population by glucose starvation, we purified the population using a lentiviral reporter system that expresses the zeocin resistance gene (Zeo^R^) under the control of the human cTnT promoter (*SI Appendix*, Fig. S2*B*) (35, 36). As a proof-of-concept, we first tested the reporter system that encodes enhanced green fluorescent protein (EGFP) and Zeo^R^ separated by a T2A self-cleaving peptide (*SI Appendix*, Fig. S2*C*). Colocalization of EGFP with cTnT immunostaining demonstrated that the reporter system provides a reliable method to obtain a pure population of cardiomyocytes (*SI Appendix*, Fig. S2*D*). For subsequent purifications, we used a cTnT reporter that expresses Zeo^R^ alone that enables purification by zeocin selection. By combining glucose starvation with zeocin selection, we capitalized on two orthogonal purification methods to ensure measurement of telomeres in the cardiomyocyte population.

We measured telomere lengths in purified iPSC-CMs from the six lines by Southern blot of telomere restriction fragments and found that telomeres in DMD cells were shortened (Fig. 3*A-F*), corroborating our previous findings in DMD iPSC-CMs using Q-FISH (12, 15). Telomere fragment resolution was achieved by running the DNA on low 5% agarose gels. We reasoned that TRF2 overexpression might suffice to prevent telomere attrition since it is a core protein in the shelterin complex that interacts directly with double-stranded telomeric DNA (Fig. 3*G*) (37, 38). TRF2 recruits the other subunits of the shelterin complex, by directly binding RAP1 and TIN2 (39). In addition, reduced TRF2 levels and telomere shortening have been observed in the hearts of patients with idiopathic and ischemic dilated cardiomyopathy (40). We transduced TRF2 on Day 10 of differentiation and found that it significantly prevented telomere attrition in all three lines of DMD iPSC-CMs twenty days later (Fig. 3*A-F*).

**Figure 3.**
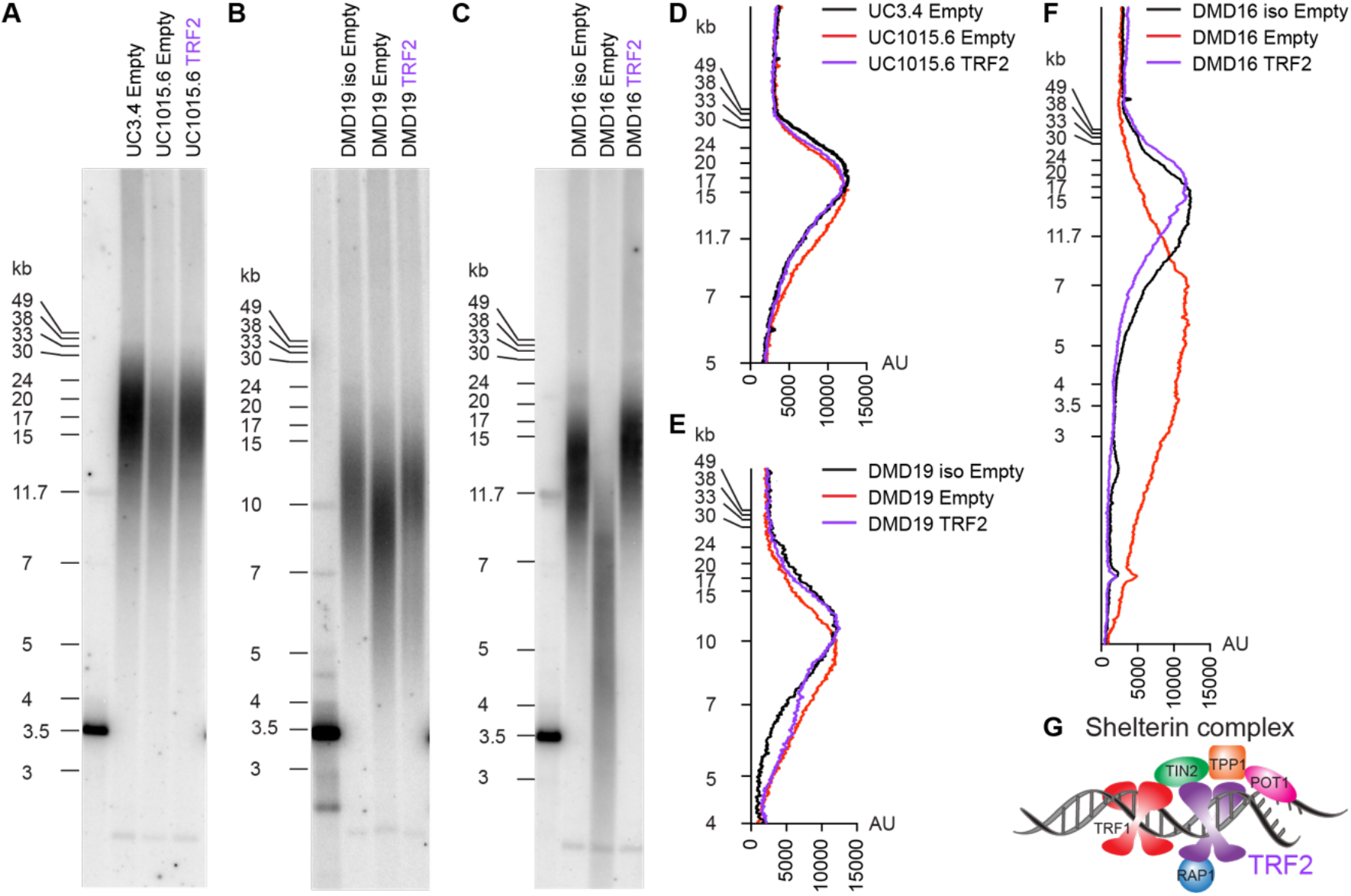
TRF2 overexpression rescues telomere attrition. Cells were transduced with TRF2 or an empty lentiviral control. Southern blot of telomere restriction fragments of Day 30 iPSC-CMs from (*A*) UC iPSC-CMs, (*B*) DMD19 iPSC-CMs, and (*C*) DMD16 iPSC-CMs. Signal distribution of telomere lengths from the Southern blot in represented as a gray value in arbitrary units for (*D*) UC iPSC-CMs, (*E*) DMD19 iPSC-CMs, and (*F*) DMD16 iPSC-CMs. (*G*) The shelterin complex is made up of six subunits. TRF1 and TRF2 directly bind telomere sequences.

As a negative control for retroviral transduction, we used an Empty retroviral control with no open reading frame. Although the magnitude of telomere attrition differed across lines, in comparison with the isogenic controls, the shortest telomeres were significantly shorter in the DMD iPSC-CMs, and these short telomeres were absent following TRF2 upregulation. These data show that protection of telomeres with TRF2 curbs telomere attrition, which is a hallmark of cardiac disease progression.

### TRF2 Attenuates the DNA Damage Response and Prolongs Cell Survival

While TRF2 protein levels were quite low in the differentiated healthy and DMD iPSC-CMs, when overexpressed, TRF2 levels were significantly elevated in all three lines of DMD iPSC-CMs (Fig. 4*A*). Because TRF2 inhibits the ataxia-telangiectasia-mutated (ATM)-mediated DNA damage response (41,42), we measured levels of phospho-CHK2, the checkpoint effector kinase that is activated in response to double-stranded breaks (43). Following TRF2 overexpression, we observed attenuated levels of phospho-CHK2 in all three lines of DMD iPSC-CMs (Fig. 4*B*). Complementary to our findings, overexpression of a dominant negative mutant of TRF2 or knocking down TRF2 with an antisense oligonucleotide resulted in CHK2 activation and telomere shortening in cardiomyocytes isolated from rats (40). We compared the survival of DMD iPSC-CMs with and without TRF2 overexpression. We found that TRF2 increased the percentage of cells that survived to Day 40 of differentiation from Day 30 (Fig. 4*C-E*). Collectively, these results show that protection of telomeres with TRF2 attenuates the DNA damage response and prevents the premature loss of cardiomyocytes, potentially delaying fibrosis and tissue stiffening in the DMD heart. This therapeutic strategy may also be relevant to other cardiomyopathies beyond those caused by dystrophin deficiency where telomere attrition is a hallmark of disease progression.

**Figure 4.**
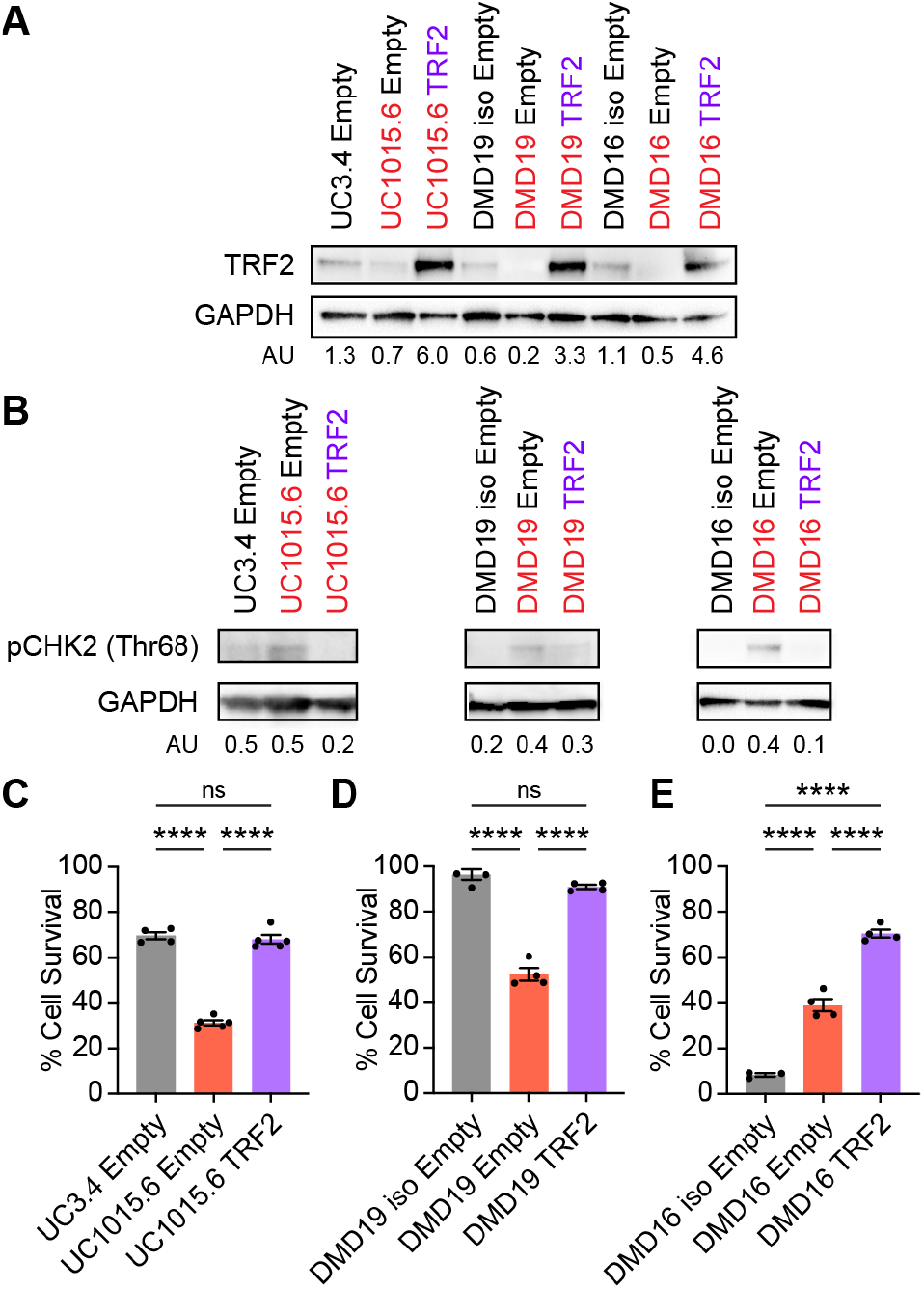
TRF2 attenuates ATM-mediated DNA damage response and prolongs cell survival. (*A*) Western blot of TRF2 levels with GAPDH as a loading control. TRF2 signal normalized to GAPDH signal in arbitrary units. (*B*) Western blot of CHK2 phosphorylated at threonine 68. Phospho-CHK2 signal normalized to GAPDH signal in arbitrary units. Percentage of cells that survived on Day 40 when compared to Day 30 of differentiation for (*C*) UC iPSC-CMs, (*D*) DMD19 iPSC-CMs, and (*E*) DMD16 iPSC-CMs. Survival was scored from 3-5 differentiation experiments. N = 375 - 12,036 cells on Day 30. Data shown are mean ± SEM. One-way ANOVA and Tukey test for post-hoc comparison was used to calculate significance. **** p < 0.0001.

### TRF2 Rescues Deficits in Cell Morphology and Improves Calcium Handling

We found that DMD iPSC-CMs exhibited decreased cell size, nuclear size, and sarcomere density compared to their healthy counterparts (Fig. *2A-L*) and sought to determine if TRF2 upregulation could ameliorate these deficits. We discovered we discovered that DMD iPSC-CMs were characterized by a diminished cell and nuclear size, both of which were improved upon TRF2 upregulation in all three DMD lines (Fig. *5A-I*). TRF2 upregulation also significantly increased the density of cTnT in UC1015.6 and DMD19 iPSC-CMs (Fig. *5J-L*). The increase in cTnT protein levels detected by immunocytochemistry were validated by western blot in the DMD19 line (*SI Appendix*, Fig. S2*F*).

**Figure 5.**
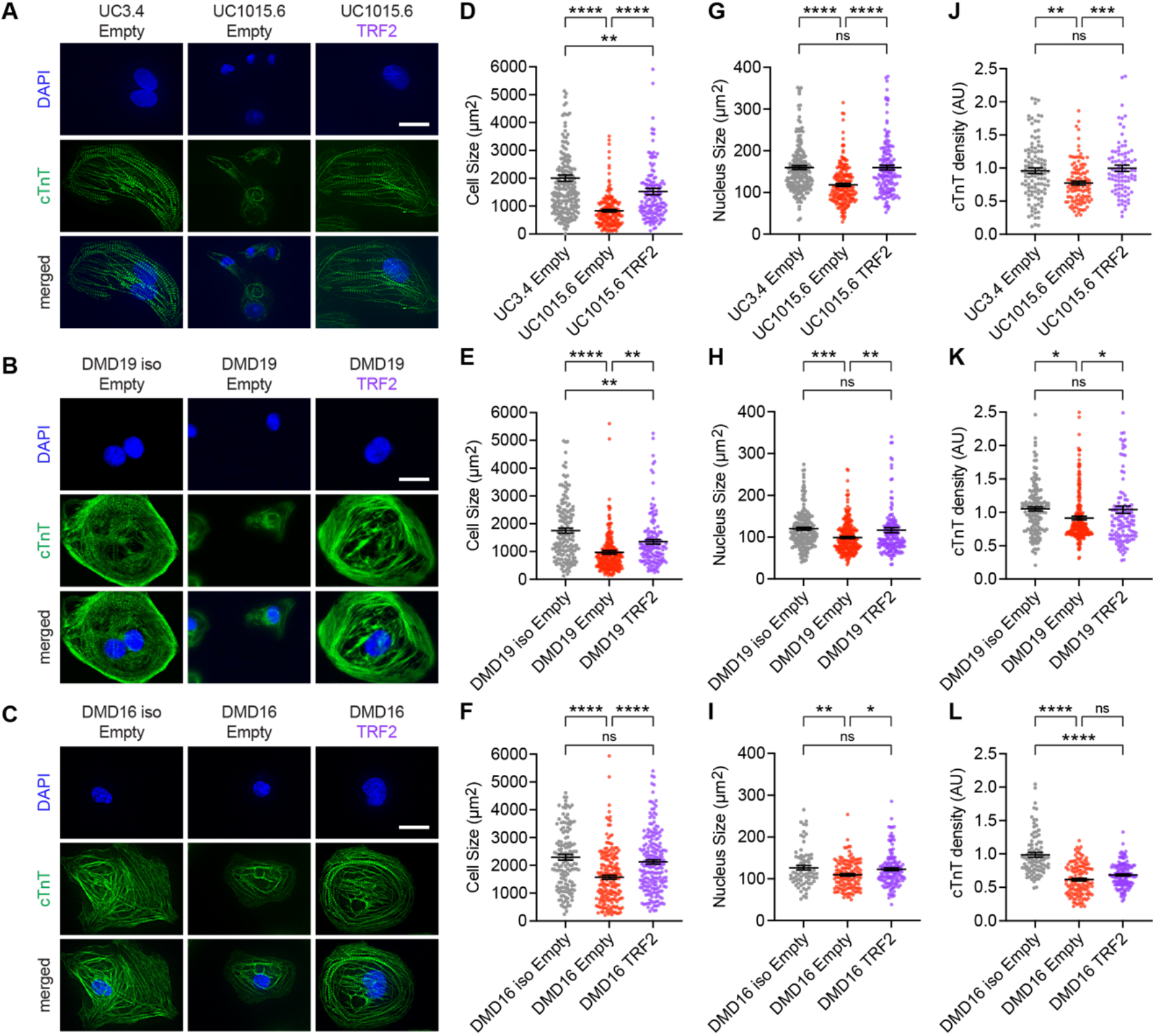
TRF2 rescues deficits in cell size, nuclear size, and sarcomere density. Cells were transduced with TRF2 or an empty lentiviral control on Day 10 and assayed on Day 30. Cardiac troponin T (cTnT) immunostaining and DAPI staining for nuclei in (*A*) UC iPSC-CMs, (*B*) DMD19 iPSC-CMs, and (*C*) DMD16 iPSC-CMs. Scale bar = 20 μm. Area of cells for (*D*) UC iPSC-CMs, (*E*) DMD19 iPSC-CMs, and (*F*) DMD16 iPSC-CMs. Nuclear size for (*G*) UC iPSC-CMs, (*H*) DMD19 iPSC-CMs, and (*I*) DMD16 iPSC-CMs. Sarcomere density as measured by cTnT signal over cell area for (*J*) UC iPSC-CMs, (*K*) DMD19 iPSC-CMs, and (*L*) DMD16 iPSC-CMs. Cells were scored from 3 differentiation experiments. N = 90-230 cells. Data shown are mean ± SEM. One-way ANOVA and Tukey test for post-hoc comparison was used to determine significance. * p < 0.05, ** p < 0.01, *** p < 0.001, and **** p < 0.0001.

In DMD cardiomyocytes, dystrophin deficiency leads to a leaky plasma membrane that causes elevated calcium levels and poor calcium handling, which correlates with arrhythmia. To evaluate the effect of TRF2 on cardiomyocyte function, we performed calcium imaging using the ratiometric calcium dye, Fura Red AM. We micropatterned iPSC-CMs at a 7:1 length:width aspect ratio on glass-bottomed wells (*SI Appendix*, Fig. S3*A*), a bioengineering method that we and others showed promotes parallel alignment of the myofibrils and improves the maturity of iPSC-CMs (15, 30, 31). TRF2 transduction restored the transient amplitude for UC1015.6 iPSC-CMs to levels comparable to the healthy isogenic control, UC3.4 iPSC-CMs, under conditions of pacing at 1 Hz, an electrical stimulation that induces contraction at one-second intervals (Fig. 5*A-B*). While other parameters of calcium handling, including time to peak, peak duration, and peak frequency, showed a partial rescue, none improved to statistically significant levels (*SI Appendix*, Fig. *S3B-M*).

We characterized the types of traces for each cell as Class I (traces with regular peaks at 1-second intervals), Class II (traces that deviate slightly from the 1-second intervals and show signs of early afterdepolarization), and Class III (traces that deviate significantly from the one-second intervals) (Fig. 5*C*). Early afterdepolarization events show calcium spikes within the decay phase of the calcium transients and can cause lethal arrhythmias (44); therefore, correction of such events would have a meaningful impact on preventing heart failure. As the iPSC-CMs are paced at 1 Hz, calcium transients should oscillate at regular 1-second intervals. Indeed, for the healthy control iPSC-CMs, the majority of cells exhibited Class I traces (Fig. *5E-G*). For the UC and DMD19 lines, DMD iPSC-CMs showed significantly higher frequencies of aberrant calcium handling of Class II and III trace types (Fig. *5E-F* and *SI Appendix*, Table S1).

Transduction with TRF2 in DMD iPSC-CMs reduced the frequency of Class II and III calcium traces (Fig. 5*E-F*); however, these reduced frequencies were not statistically significant. Based on the variance of the peak duration and variance of the amplitude, an arrhythmia score was calculated for each cell. The scoring system was weighted 90% on peak duration variance and 10% on amplitude variance on a ten-point scale (Fig. 6*D*). There was a significant difference in arrhythmia scores between DMD and healthy isogenic controls for the UC and DMD19 lines, and TRF2 treatment improved the score, but this change was not statistically significant (Fig. *6H-J*). These results collectively demonstrate that TRF2 upregulation not only confers telomere protection and prevents telomere attrition but provides morphological advantages that impart functional improvements to DMD iPSC-CMs.

**Figure 6.**
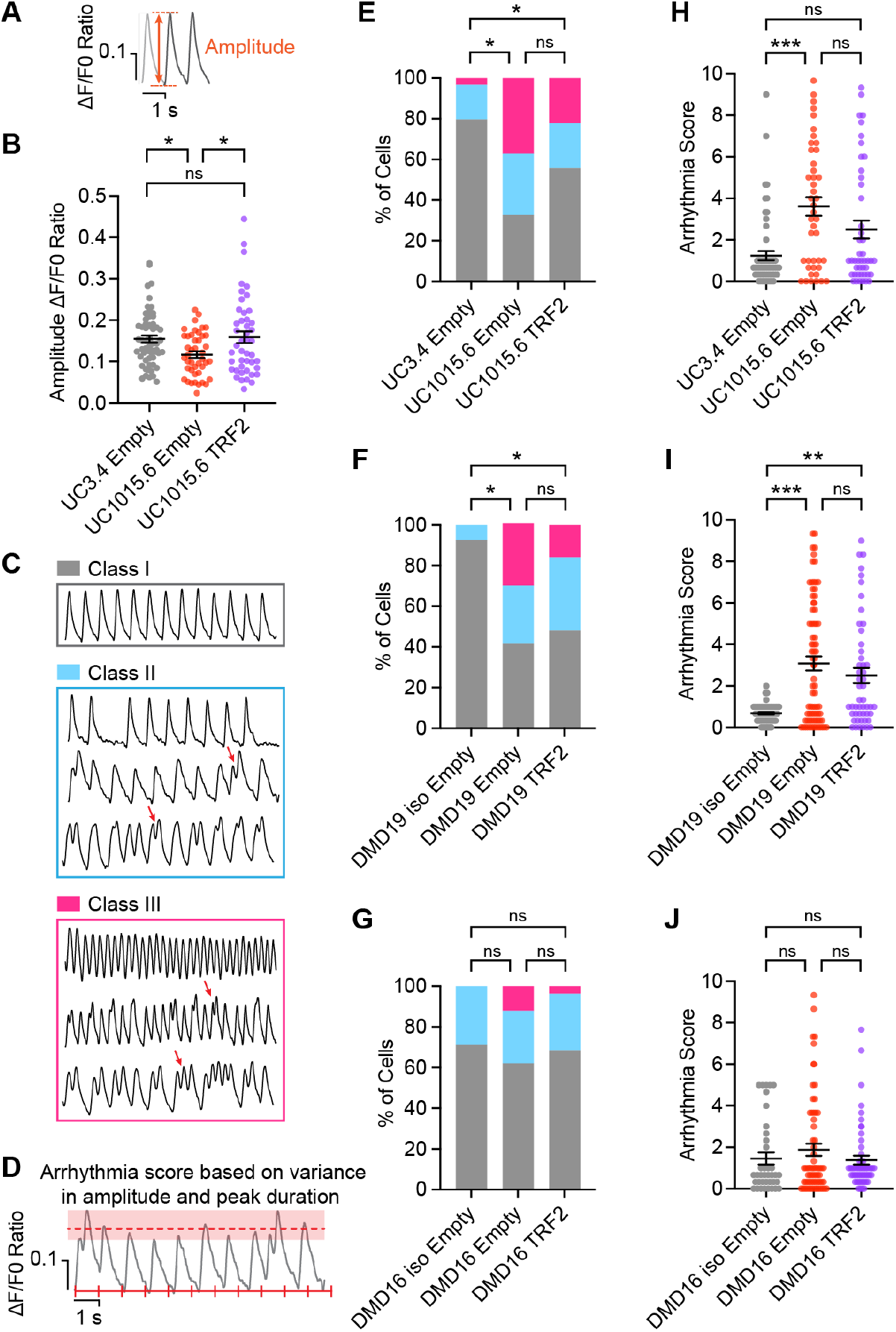
TRF2 partially rescues aberrant calcium handling in DMD iPSC-CMs. Cells were transduced with TRF2 or an empty retroviral control on Day 10 and assayed on Day 30. (*A*) Amplitude is the difference between the maximum and minimum calcium ratio. (*B*) Amplitude for UC iPSC-CMs. (*C*) Class I – traces with regular peaks at 1-second intervals, Class II – traces that deviate slightly from the 1-second intervals and show signs of early afterdepolarization, and Class III – traces that deviate significantly from the 1-second intervals. Examples of early afterdepolarization events are indicated by the red arrows. (*D*) Arrhythmia scores were determined on a 10-point scale based on variance of peak duration (90%) and variance of amplitude (10%). Calcium traces categorized into three classes for (*E*) UC iPSC-CMs, (*F*) DMD19 iPSC-CMs, and (*G*) DMD16 iPSC-CMs. Pearson’s chi-square test followed by z-test with Bonferroni correction was used for pairwise comparisons. \**p* < 0.05. See *SI Appendix*, Table S1 for z-scores. Arrhythmia score for (*H*) UC iPSC-CMs, (*I*) DMD19 iPSC-CMs, and (*J*) DMD16 iPSC-CMs. One-way ANOVA and Kruskal-Wallis test for post-hoc comparison was used to determine significance. ** p < 0.01 and *** p < 0.001. Calcium imaging from 4-6 differentiation experiments performed on 38-77 cells per condition. Cells were paced at 1 Hz. Data shown are mean ± SEM.

## Discussion

This study shows that treatment with TRF2 prevents telomere attrition, restores morphological deficits observed in DMD iPSC-CMs, and prolongs cells survival. Our previous findings in mice revealed that shortening the telomeres of dystrophin-deficient *mdx^4cv^* mice by breeding *mdx*^4*cv*^ to *mTR* knockout mice provides a “humanized” model to study the severe phenotypes of Duchenne muscular dystrophy. The resulting *mdx*^4*cv*^/*mTR*^*G*2^ mice exhibit severe skeletal muscle degeneration, kyphosis, atrophied diaphragms, dilated cardiomyopathy, and die prematurely (8–10). In histological sections of human heart tissue, telomeres are shorter in DMD patient cardiomyocytes compared to age-matched healthy controls (12), underscoring the role of telomere attrition in the etiology of dilated cardiomyopathy. To gain insights into the role of telomere shortening in the demise of DMD cardiomyocytes, we used human iPSCs as a disease model. Previously, we demonstrated that telomere attrition, poor calcium handling, and contractile dysfunction are recapitulated in DMD iPSC-CMs (15). Here we validate telomere attrition by Southern blotting of telomere restriction fragments, the gold standard for telomere length measurements, which reinforces our previous finding obtained by Q-FISH and monochrome multiplex quantitative PCR (12, 15). Importantly, we demonstrate that DMD phenotypes can be rescued, providing novel mechanistic insights and a potential therapeutic intervention.

We hypothesized that telomere attrition plays a role in the cardiac failure seen in DMD patients and sought to rescue the cardiomyocyte phenotypes observed in vitro in DMD iPSC-CMs by protecting telomeres. Previously, we reported that subunits of the shelterin complex are expressed at lower levels in DMD iPSC-CM compared to healthy controls (15). TRF2, a core subunit of the shelterin complex, can repress the ATM-mediated DNA damage response (41,42), inhibit binding of poly(ADP-ribose) polymerase 1 (PARP1) to telomeres (45), and prevent hyperresection of telomeres by exonucleases (46). Here, we show that TRF2 upregulation not only prevents telomere attrition but ameliorates morphological deficits and prolongs cell survival in DMD iPSC-CMs. These results suggest that telomere attrition contributes to the pathogenesis and demise of cardiomyocytes and that preservation of telomeres improves the differentiated state through regulation of cell growth and maturation. While we observed signs of improving calcium handling in the DMD iPSC-CMs, functional rescue was incomplete because full-length dystrophin is still missing in the DMD iPSC-CMs. Notably, the rescue of telomere attrition and enhanced cell survival were observed regardless of mutation underscoring the critical role of telomeres in the etiology of DMD. The strength of our findings is underscored by the use of three DMD lines and three isogenic controls where the dystrophin mutations are corrected.

Protection of telomeres by TRF2 can potentially be a therapeutic strategy to ameliorate other cardiovascular diseases where we and others have observed telomere attrition (12, 40, 47). Our findings of preventing telomere attrition in DMD fit well with prior pioneering research showing that telomerase gene therapy reverses aplastic anemia and pulmonary fibrosis in mice through telomere elongation (48, 49). Our observations of TRF2 in DMD are also in line with the groundbreaking discovery that another subunit of the shelterin complex, TRF1, prolongs the lifespan of mice and protect against anemia and metabolic dysregulation during aging (50). Whether TRF2 overexpression rescues telomere attrition in iPSC-CMs harboring mutations in other proteins that make up the contractile apparatus remains to be elucidated. TRF2 is amenable to gene therapy as the coding sequence can easily fit into adeno-associated virus (51). While TRF2 does not replace the function of dystrophin, we hypothesize that delivery of TRF2 can delay the premature loss of cardiomyocytes and prevent the fibrosis that accelerates the onset of dilated cardiomyopathy. Our results demonstrate that TRF2 provides an advantage to DMD iPSC-CMs irrespective of the type of mutation and highlights protection of telomeres as a novel therapeutic target for other dilated and hypertrophic cardiomyopathies that manifest telomere attrition (12).

## Materials and Methods

### Differentiation to Cardiomyocytes

For maintenance of iPSCs, cells were seeded on Matrigel (Corning 356231)-coated plates and were cultured in Nutristem media (Reprocell 01-0005) with daily media change. For differentiation to cardiomyocytes, iPSCs were cultured to 70-90% confluency and cultured in RPMI-1640 supplemented with 1X B27 minus insulin and 4-6 μM CHIR-99021 (Selleck S2924). Two days later, cells were refreshed with RPMI-1640 supplemented with 1X B27 minus insulin and 5 μM IWR-1 (Sigma-Aldrich I0161). Two days later, cells were refreshed in RPMI-1640 supplemented with 1X B27 minus insulin. Two days later, cells were refreshed in RPMI-1640 supplemented with 1X B27 and maintained for 4 days with a media change every other day. From day 10 of differentiation onward, iPSC-CMs were maintained in RPMI-1640 minus glucose supplemented with 1X B27 and 4 mM lactate (Sigma-Aldrich L7900) with media change every other day. On Day 10, iPSC-CMs were transduced with TRF2 or an empty control retrovirus with 8 μg/mL polybrene. Media was changed every two days. On Day 15, cells were harvested with Accutase:TrypLE (1:1 v/v) and replated on Matrigel-coated plates in RPMI-1640 minus glucose supplemented with 1X B27, 4 mM lactate, 5% KSR, and 10 μM Y-27632. On Day 16, iPSC-CMs were transduced with TNNT2-zeo. On Day 18-19, cardiomyocytes were selected for resistance to zeocin (20 μg/mL). All assays were performed on Day 30 of differentiation.

### Cloning Viral Vectors

Human TRF2 under control of a CMV promoter was expressed from pLPC-TRF2, a gift from Titia de Lange (Addgene plasmid #18002). A short cassette without an open reading frame flanked by HindIII and EcoRI sites was cloned into pLPC-TRF2 to generate the pLPC-Empty plasmid. TroponinT-GCaMP5-Zeo was a gift from John Gearhart (Addgene plasmid #46027). GCaMP5 was substituted for EGFP to generate TroponinT-EGFP-Zeo. A vector expressing the zeocin resistance gene only, TroponinT-Zeo, was created by cloning the zeocin resistance gene downstream of the promoter.

### Virus Production

Retrovirus was produced in Phoenix Amphotropic cells by transfection with FUGENE 6 (Promega E2692) of pLPC-TRF2 or pLPC-Empty. Lentivirus was produced in HEK293T cells by transfection with FUGENE 6 of TNNT2-Zeo or TNNT2-GFP-Zeo, psPAX2 packaging, and pMD2.G envelope plasmids. Media containing virus as harvested 78 hours post-transfection. Retrovirus or lentivirus was concentrated using Lenti-X Concentrator (Takara Bio 631231) using the manufacturer’s protocol.

### Immunofluorescence

For immunostaining, iPSCs or Day 27 iPSC-CMs were seeded onto glass slides coated with Matrigel. Three days later, cells were fixed in 4% paraformaldehyde in PBS for 15 minutes. Cells were blocked and permeabilized in buffer (20% goat serum, 0.3% Triton X-100 in PBS) for 1 hour at room temperature. Cells were incubated in primary antibody for 2 hours at room temperature or overnight at 4°C, then washed 3 × 5 minutes in PBS. Cells were stained with the following antibodies: cardiac troponin T (Thermo Fisher MA5-12960, 1:200 dilution), dystrophin (Abcam 15277; 1:200 dilution), dystrophin (DSHB MANEX1A; 1:200 dilution), OCT4 (Millipore MAB4419; 1:100 dilution), SOX2 (BD Pharmingen 561593; 1:100 dilution), or TRA-1-60 (ESI-BIO ST11016; 1:100 dilution). Cells were incubated with secondary antibody for 1 hour at room temperature, then washed 3 x 5 minutes in PBS, with the final two washes containing 3 nM DAPI. Coverslips were mounted with Fluoromount-G (Southern Biotech 0100-01). Images were acquired on Everest deconvolution workstation (Intelligent Imaging Innovations) equipped with a Zeiss AxioImager Z1 microscope and a CoolSnapHQ cooled CCD camera (Roper Scientific). A 20X NA0.5 Plan-Apochromat objective lens (Zeiss 420762-9800) or 63X NA1.4 Plan-Apochromat objective lens (Zeiss 420780-9900) were used to capture micrographs. Deconvolution of 3D micrographs taken at 63X were processed with Microvolution. Size and fluorescence instensities were analyzed using ImageJ.

### Western Blot

Cells were flash frozen and lysed in lysis buffer (50 mM Tris-HCl, pH 7.5, 150 mM NaCl, 1% Triton X-100, 0.1% Na deoxycholate). 25 μg of total protein was loaded on a NuPAGE Bis-Tris gel using 1X NuPAGE buffer and beta-mercaptoethanol (2.5% final concentration). Gel was run in MOPS buffer for 1-2 hours at 100V. A wet transfer to nitrocellulose membrane was done using NuPAGE Transfer Buffer at 4°C. Membrane was blocked in 5% milk for 1 hour, then in primary antibody (TRF2 Novus NB100-56506 at 1:500; phosphor-CHK2 Thr68 Cell Signaling C13C1 at 1:1000; GAPDH Cell Signaling D16H11 at 1:1000) overnight 4°C, and Goat anti Mouse IgG-HRP at 1:4000 (Thermo Fisher Scientific 62-652-0) or Goat anti Rabbit IgG-HRP at 1:2000 secondary antibody for 1 hour. Between incubations membrane was washed 3 × 10 minutes in Tris Buffered Saline with 0.1% Tween-20. Membrane was imaged by chemiluminescence (Nacalai Tesque 07880) on a Bio-Rad ChemiDoc XRS System.

### Telomere Restriction Fragment Southern Blot

Cells were flash frozen and lysed in lysis buffer (100 mM Tris HCl, 5 mM EDTA, 200 mM NaCl, 0.2% SDS, and 60 μg/mL Proteinase K). DNA was extracted with phenol:chloroform:isoamyl alcohol. The aqueous layer was treated with 240 μg/mL RNase A for 37°C for 1 hour before extraction with chloroform. DNA was ethanol precipitated with 75 μg/mL sodium acetate. Tris-EDTA buffer pH 8.0 was added to precipitated DNA and incubated overnight at 37°C. DNA integrity was verified on a 1% agarose 0.5X TBE gel. DNA was digested with MboI and AluI for 20 hours at 37°C. Digested DNA was run on a 0.8% agarose 0.5X TBE gel to verify adequate digestion. 3-5 μg of DNA was loaded on a 0.5% agarose 1X TBE gel at 85V for 24 hours at 4°C. Gel was stained with ethidium bromide afterward to ensure proper migration. Gel was washed in depurination buffer (0.25 N HCl) for 30 minutes, denaturing buffer (1.5 M NaCl, 0.5 M NaOH) for 1 hour, and neuralization buffer (1.5 M NaCl, 1 M Tris-HCl pH 7.4) for 1 hour at room temperature. DNA was transferred from the gel to a Hybond N membrane (GE Healthcare RPN203N) by capillary transfer. DNA was crosslinked to the membrane using the Strategene UV Stratalinker 1800. P32 end-labeled GGGTTA3 probe was hybridized to the membrane in Rapid-Hyb Buffer (GE Healthcare RPN1635) at 42°C for 2 hours. Membrane was washed in 2X SSC with 0.1% SDS for 20 minutes at 42°C. Membrane was washed in 1X SSC with 0.1% SDS for 2 × 15 minutes at 42°C. Membranes were air dried for 1 hour, then exposed on an intensifier screen for 24-48 hours. Images were scanned using a phosphor imager (GE Typhoon).

### Calcium Imaging

For ratiometric calcium imaging, we imaged Fura Red, AM (ThermoFisher F3012) on the Zeiss AxioObserver inverted microscope and a Photometrics PRIME 95B sCMOS camera, using a custom-made filter cube with dual band excitation filter (430/24 nm and 470/40 nm), dichronic mirror 505 nm and emission filter 670/50 nm (Chroma), and a Colibri 7 Type R fast switching LED illumination light source. Pseudo-simultaneous videos were recorded in a longitudinally-cropped area along the cardiomyocyte main axis using excitation wavelengths of 471 nm and 435 nm and emission wavelengths of 670 nm and 660 nm at 115 fps/channel for 14 sec. Cells were paced at 1 Hz and maintained at 37°C with 5% CO2 in a humidified chamber. Since Fura Red is a calcium quenching dye, ratiometric signals were calculated as signal at 471 nm/signal at 435 nm. Videos were analyzed using a custom MATLAB script.

### Cell Viability Assay

Calcein Blue, AM (Thermo Fisher C34853) was used to measure viability at Day 30 and Day 40 of differentiation. Cells were incubated with 5 μM Calcein Blue for 15 minutes at 37°C. After dye uptake, cells were imaged in RPMI 1640 without phenol red and glucose supplemented with 1X B27 and 4 mM lactate. Images of the same field were captured with a 10X objective on a Keyence microscope. The percentage of cells that survived was calculated by dividing the number of cells fluorescently labeled on Day 40 by the number on Day 30.

## Acknowledgments

We are grateful to David L. Mack, Martin K. Childers, and Chris Denning for providing the iPSC lines. We acknowledge Garry P. Nolan for providing the Phoenix Amphotropic cells. This work was supported by the American Heart Association (18POST33960526 to A.E. and 17CSA33590101 to H.M.B.), Stanford School of Medicine Dean’s Postdoctoral Fellowship (A.E.), Stanford Translational Research and Applied Medicine Pilot Grant (A.E.), the Stanford Bio-X Summer Undergraduate Research Program (F.B.), a Major Grant from the Stanford University Vice Provost for Undergraduate Education (F.B.), Tobacco-Related Disease Research Program Fellowship 26FT-0029 (J.Z.Z.), the National Institutes of Health HL159340 (H.M.B.), the Keck Foundation (H.M.B.), the Baxter Foundation (H.M.B.), and the Li Ka Shing Foundation (H.M.B.). For statistical analysis, we consulted the Stanford’s Data Studio, supported by the National Center for Advancing Translational Sciences of the National Institutes of Health under Award Number UL1TR003142.

## Author Contributions

A.E. and H.M.B. designed research; A.E., K.K., and A.F.G.G. performed research; A.E., K.K., S.I.T.B., and A.F.G.G. analyzed data; F.B. contributed new analytic tools; and A.E. and H.M.B. wrote the paper.

## Supplementary Information Appendix

**Figure S1.**
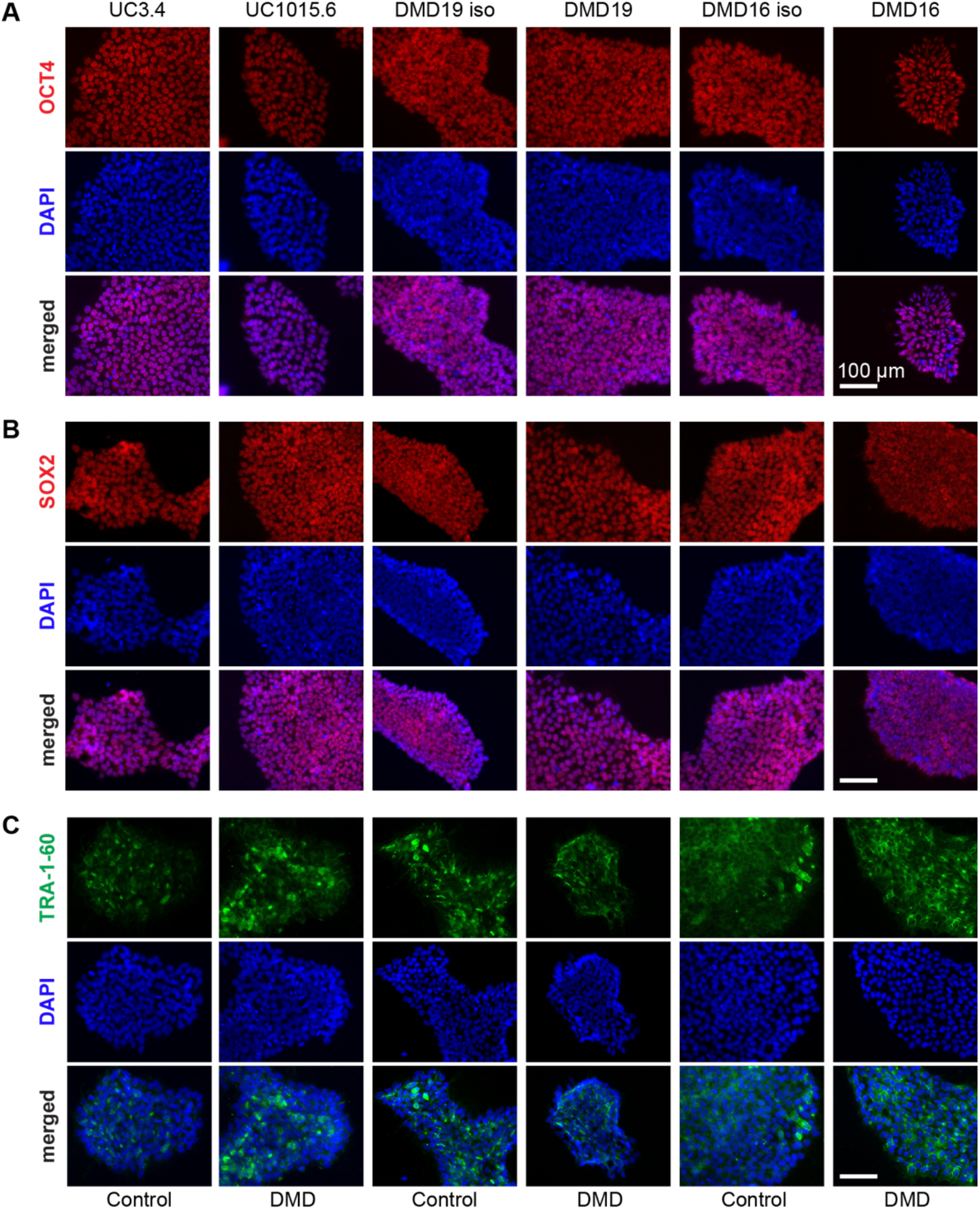
Markers of pluripotency expressed by iPSCs. Immunostaining for (*A*) OCT4, (*B*) SOX2, and (*C*) TRA-1-60. Scale bar = 100 μm.

**Figure S2.**
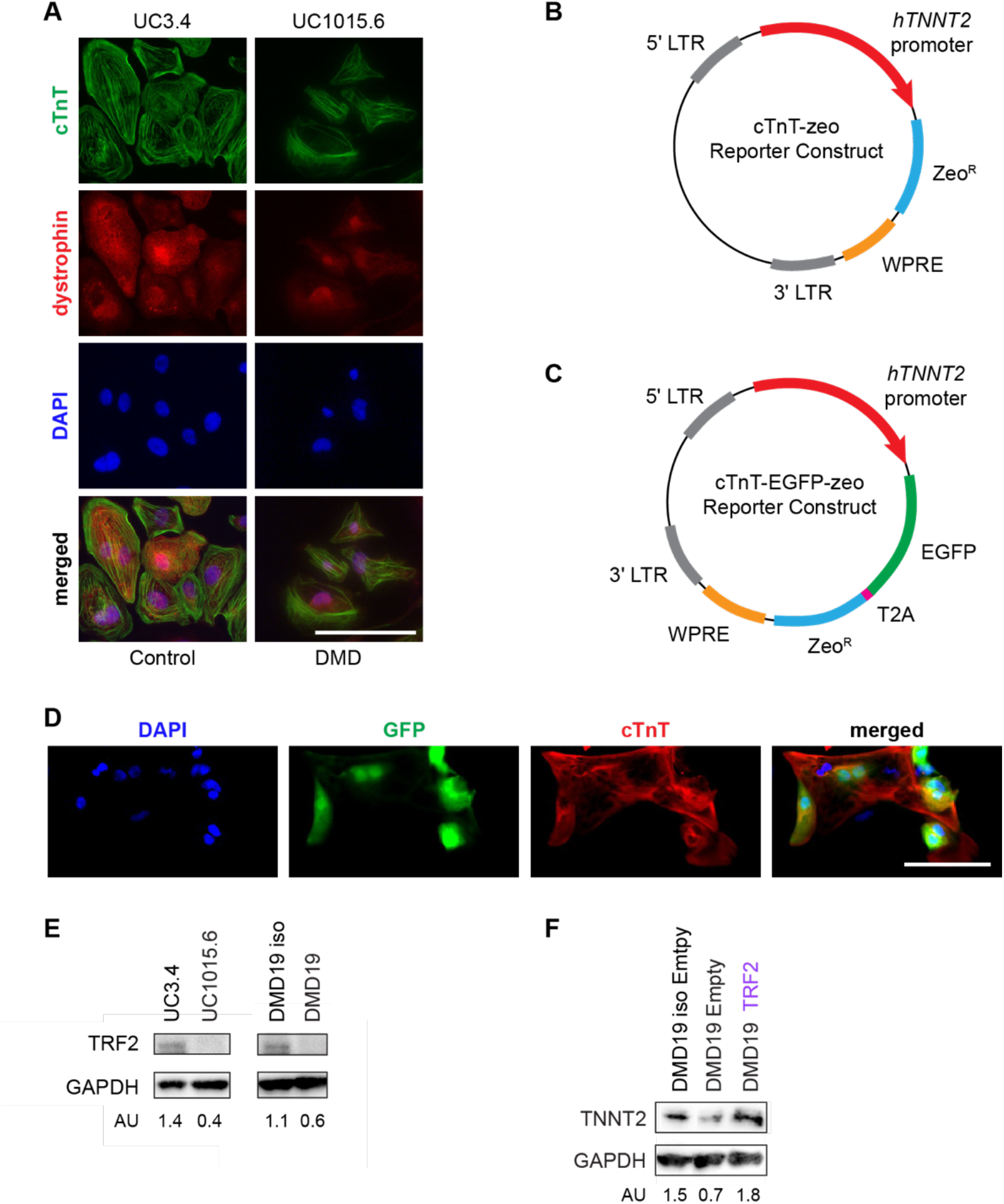
Cardiomyocyte markers in iPSC-CMs. (*A*) Immunostaining for dystrophin in Day 30 iPSC-CMs using the ab15277 antibody that recognizes the C-terminus. Cardiac troponin T is a cardiac-specific marker. DAPI marks nuclei. Scale bar = 100 μm. (*B*) Cardiac troponin T reporter construct. The human troponin T promoter drives expression of the bicistronic cassette that expresses EGFP and the zeocin resistance gene separated by a T2A self-cleaving peptide. (*C*) Non-fluorescent cardiac troponin T reporter construct. The human troponin T promoter drives expression of the zeocin resistance gene only. (*D*) EGFP reporter in iPSC-CMs immunostained for cardiac troponin T. DAPI marks nuclei. Scale bar = 100 μm. (*E*) Western blot of TRF2 levels with GAPDH as a loading control in Day 30 iPSC-CMs. TRF2 signal normalized to GAPDH signal in arbitrary units. (*F*) Western blot of cardiac troponin T (TNNT2) levels with GAPDH as a loading control in Day 30 iPSC-CMs. TNNT2 signal normalized to GAPDH signal in arbitrary units.

**Figure S3.**
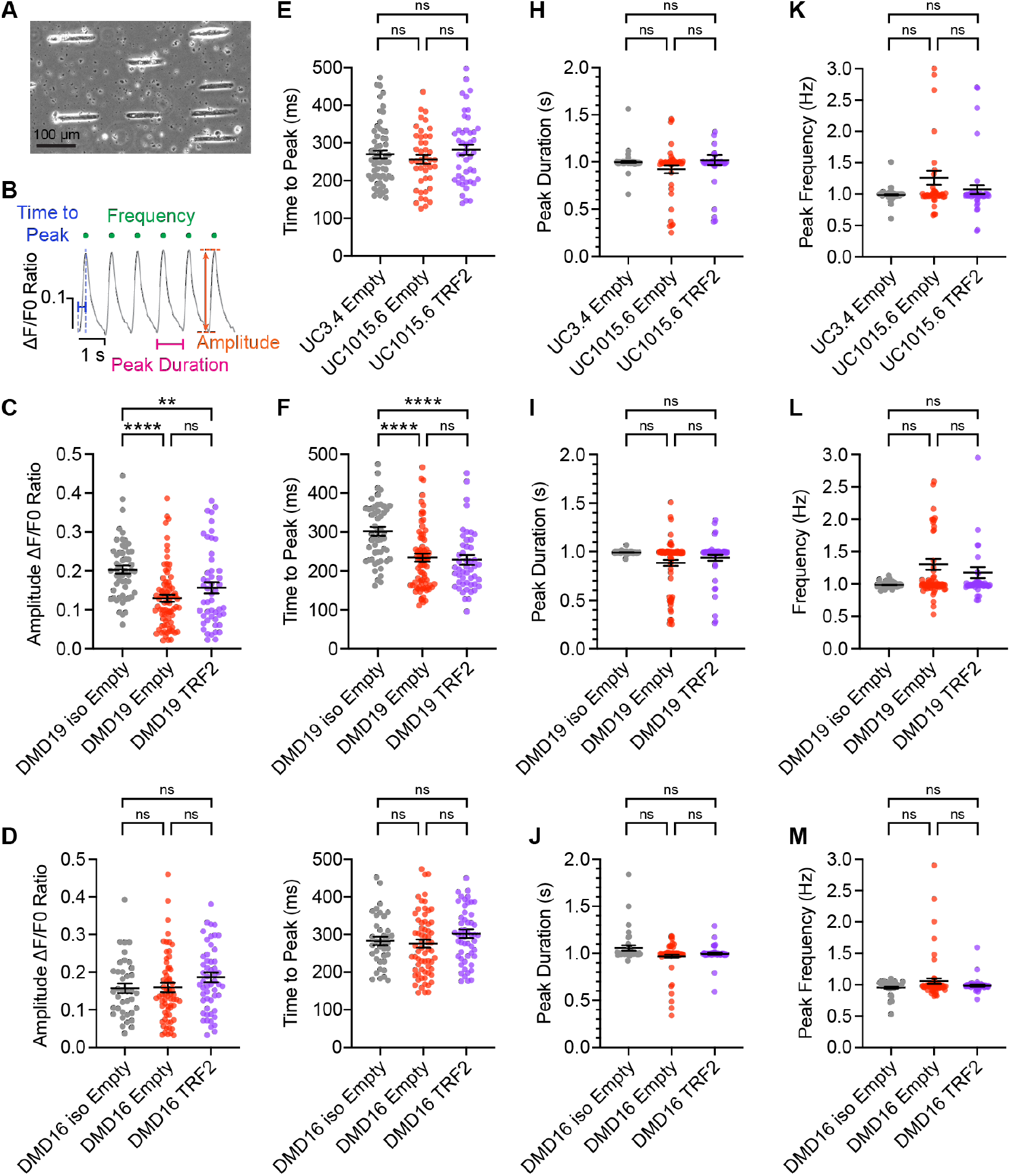
Effects of TRF2 on calcium handling of Day 30 iPSC-CMs paced at 1 Hz. Cells were transduced with TRF2 or an empty retroviral control. (*A*) Micropatterning of iPSC-CMs to a 7:1 length:width aspect ratio promotes maturation by parallel alignment of the myofibrils and adoption of a physiologically-relevant morphology. (*B*) Parameters of calcium handling. Amplitude is the difference between the maximum and minimum calcium ratio. Time to peak is the duration of time from minimum to maximum. Peak duration is the duration of time from minimum to the next minimum. Peak frequency is the rate of oscillations per second. 1 Hertz (Hz) is equal to 1 cycle per second. Amplitude for (*C*) DMD19 iPSC-CMs and (*D*) DMD16 iPSC-CMs. Time to peak for (*E*) UC iPSC-CMs, (*F*) DMD19 iPSC-CMs, and (*G*) DMD16 iPSC-CMs. Peak duration for (*H*) UC iPSC-CMs, (*I*) DMD19 iPSC-CMs, and (*J*) DMD16 iPSC-CMs. Peak frequency for (*K*) UC iPSC-CMs, (*L*) DMD19 iPSC-CMs, and (*M*) DMD16 iPSC-CMs. Cells were scored from 4-6 differentiation experiments. Calcium imaging was performed on 38-77 cells per condition. Data shown are mean ± SEM. One-way ANOVA was used to determine significance. For pair-wise comparisons, Tukey test was used for normally distributed data and Kruskal-Wallis for data not normally distributed. * p < 0.05, ** p < 0.01, and **** p < 0.0001.

**Table S1.**
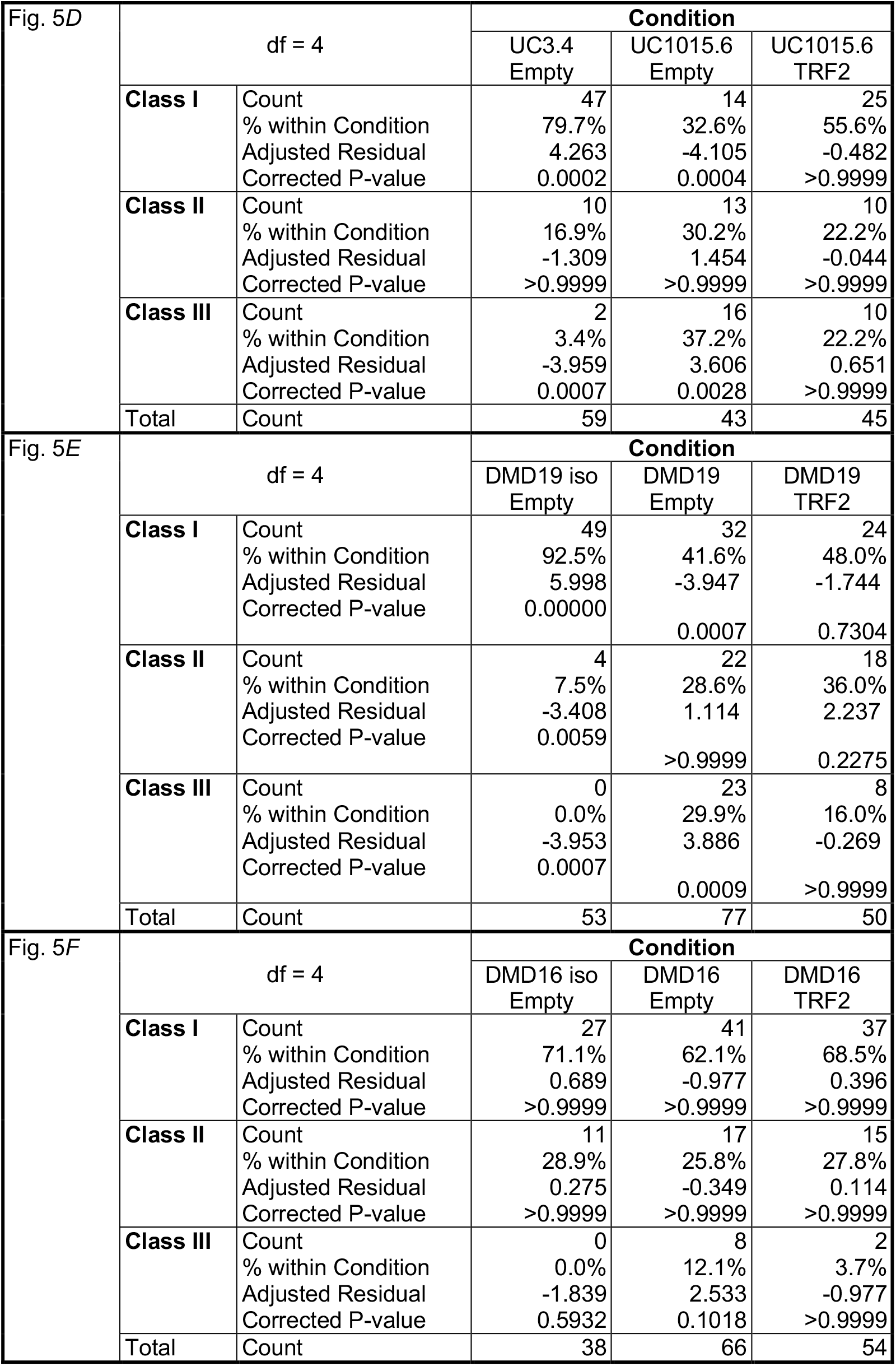
Chi-squared test for Figure *5D-F*.

